# Stem Cell Secretomes from Surgical Waste: A Novel Approach to Cancer Therapy – An In Vitro Study

**DOI:** 10.1101/2025.06.05.656255

**Authors:** Vinod Kumar Verma, Syed Sultan Beevi, Nagesh Kishan Panchal, Kondapuram Sreekarani, Radhika Chowdary Darapuneni

## Abstract

**Introduction:** Cancer is the second leading cause of mortality in India, with conventional chemotherapy often limited by drug resistance, off-target toxicity, and metastasis promotion. These treatments indiscriminately attack rapidly dividing cells, including healthy tissues and immune cells, leading to severe side effects and weakened antitumor immunity. Hence, alternative strategies are needed to selectively target cancer cells while minimizing collateral damage.

**Methods:** Mesenchymal stem cells (MSCs) were isolated, cultured up to passage 4 (P4), and characterized via flow cytometry and immunocytochemistry for stemness markers. Their differentiation potential was validated using adipogenic and chondrogenic lineage assays. The MSC-derived secretome, collected under hypoxic conditions, was analyzed for its cytotoxic effects on cancer cells using MTT, TMRM staining, and scratch assays. NanoLC-MS/MS profiling and STRING-DB analysis identified key proteins involved in apoptosis, tumor suppression, and DNA damage response.

**Results:** P2 secretomes exhibited the highest protein concentration and strongest cytotoxic effects. UCSC-derived secretomes showed superior anti-cancer activity, significantly reducing cancer cell viability, mitochondrial membrane potential, and migration. Dose-dependent studies revealed up to 95% cancer cell death with UCSC secretome, while ADSC secretome achieved 80%. In contrast, HEK293-derived secretomes had minimal effects. LC-MS profiling identified tumor-suppressive and apoptosis-inducing proteins, with unique protein signatures distinguishing UCSC and ADSC secretomes. STRING analysis mapped key regulatory pathways in tumor inhibition.

**Conclusion:** MSC-derived secretomes, particularly from UCSCs, effectively target aggressive breast cancer cells while sparing normal cells, offering a promising alternative to conventional therapies. Future research should optimize secretome formulations, identify key bioactive components, and validate their efficacy in in vivo models and clinical trials to advance this therapy toward clinical applications.

## 1. Introduction

Cancer is the second most common cause of mortality in India, accounting for a significant number of deaths annually (1). Achieving disease-free survival remains a challenge due to the failure of chemotherapeutic drugs, driven by drug resistance, inherent limitations, and metastatic progression (2). Chemotherapeutic drugs, being cell-cycle-specific, indiscriminately target all rapidly dividing cells, including non-cancerous ones (3). Moreover, chemotherapy impair the host’s immune cells (white blood cells), thereby reducing the immune system’s antitumor response (4).

Thus, tumor growth should be managed in a way that minimizes collateral damage while simultaneously strengthening the antitumor immune response. In this context, utilizing the potential of the stem cell secretome to control tumor progression and enhance immune responses presents an attractive strategy for tumor therapy. Mesenchymal stem cells (MSCs) are known to secrete a plethora of bioactive molecules such as growth factors, cytokines, chemokines, and extracellular vesicles (EVs) in response to their surrounding niche (5). The secretome exhibits properties such as immunosuppression, angiogenesis, anti-inflammation, anti-apoptosis, and anti-fibrosis (6).

Studies have demonstrated the therapeutic effects of the stem cell secretome on oral squamous cell carcinoma (7), neuroblastoma (8), and drug-resistant triple-negative breast cancer (9). Bone marrow-derived mesenchymal stem cells (BMSCs) have been shown to inhibit the proliferation, migration, invasion, and apoptosis of glioma U251 cells (10). Additionally, umbilical cord-derived mesenchymal stem cell-conditioned medium has been found to reduce the growth of PC3 and LNCaP cells without affecting non-malignant epithelial prostate cells (11). Notably, MSC secretomes have been observed to downregulate AKT and PI3K signaling while upregulating p53 expression (12). Furthermore, MSC secretomes contain high levels of tumor-inhibiting factors, including TNF superfamily member 14, Fms-related tyrosine kinase 3 ligand, CXCL10, and latency-associated protein (13). Another study demonstrated that Wharton’s jelly-derived MSC (WJ-MSC) secretome is non-tumorigenic and does not induce resistance to doxorubicin in lung cancer cells (A549) (13).

MSCs have been isolated from various sources, with bone marrow being the most reliable for clinical applications. However, adipose tissue and umbilical cord tissue are increasingly considered excellent sources of MSCs due to their non-invasive collection methods, easy availability, and status as medical waste, typically discarded after procedures. This study aims to explore the potential of discarded tissues as sources of MSCs and the secretome derived from them to inhibit tumor cell proliferation, growth, and invasion without harming non-cancerous cells in vitro. Since more than 80% of the therapeutic effects of stem cells occur through their paracrine mode of action via bioactive molecule secretion, the secretome offers a promising avenue for developing advanced cancer therapies.

## 2. Materials and Methods

### 2.1 Isolation of Umbilical Cord-Derived Mesenchymal Stem Cells

Following normal childbirth, fresh human umbilical cord tissue (1–2 inches in size) was collected with informed consent and transported in sterile culture medium on ice. The umbilical cord tissues were thoroughly cleaned with phosphate-buffered saline (PBS), cut into several small pieces (2–3 cm in size, 10–15 explants), and seeded into 90 mm Petri dishes. These were placed in a CO□ incubator at 37 °C with 5% CO□ for 30 minutes to facilitate attachment. Complete medium (F12) supplemented with 20% FBS, 0.1% FGF, and 1% antibiotic was added, and the cultures were incubated for five days without disturbance. The medium was replenished every alternate day, and cell growth from the explants was monitored. Primary cells (P0) were trypsinized at 70–80% confluency using trypsin-EDTA and further cultured up to passage 4 (P4).

### 2.2 Isolation of Adipose-Derived Mesenchymal Stem Cells

Adipose tissue left over from liposuction surgeries of healthy individuals was collected in sterile carrier media on ice. The tissue was cleaned with sterile PBS, finely chopped using a scalpel, and subjected to enzymatic digestion with 0.1% collagenase type III for 30 minutes in a CO□ incubator at 37 °C and 5% CO□. Culture media with 10% FBS was added to the digested tissue, which was then filtered using a cell sieve. The filtered cells were centrifuged for 15 minutes, and the resulting cell pellet was resuspended in complete media (F12). The cells were cultured up to passage 4 (P4).

### 2.3 Morphological evaluation of stem cell

Mesenchymal stem cells isolated from adipose and umbilical cord tissue were subjected to evaluate the morphological characteristic using Geimsa staining (Sigma Aldrich). Dissolved 3.8 g of Giemsa powder in 250 ml of methanol and gradually add 250 ml of glycerin to the solution while stirring. After filter the satin was used as working solution for stem cell staining. The overnight grown stem cells were exposed t0 Geimsa solution (2.5%) for 20 to 30 min. After through wash image were mounted using DPX motuning media and imaged using an Olympus fluorescence microscope (BX53) with a 40X objective.

#### MSC Doubling Time

Mesenchymal stem cells (4 dishes, each with 5000 cells/dish) were seeded and cultured for periods of 48, 72, 96, and 120 hours. At each time point, the cells were trypsinized and counted to calculate doubling time using the exponential phase of a growth curve formula, as determined by an online doubling time calculator (http://www.doubling-time.com/compute.php).

### 2.4 Characterization of Mesenchymal Stem Cells

MSCs (1 × 10□ cells from P2 passage) were characterized for stemness markers using positive markers (CD90, CD105, CD73, FITC-labeled) and negative markers (CD34, CD45, and HLA-DR, PE-labeled). Flow cytometry antibody labeling and analysis were performed following the user manual of the stem cell characterization kit (company name). IgG isotype controls were used for normalization and gating purposes.

Immunocytochemistry was conducted by seeding MSCs (1 × 10□ cells) on 18 mm glass coverslips, followed by overnight growth. The cells were labeled with primary antibodies for positive markers (CD90, CD105, CD73) and negative markers (CD34, CD45, and HLA-DR) and stained with Alexa 488-labeled secondary antibodies. After washing, coverslips were mounted with glycerol as the mounting medium. Images were captured using an Olympus fluorescence microscope (BX53) with a 40X objective. Secondary antibody and isotype controls were used to normalize fluorescence gain. Captured images were processed using cellSens software.

### 2.5 MSC Lineage Differentiation

### 2.6 Adipogenic Differentiation

MSCs were seeded on 22 mm coverslips at a density of 2 × 10□ cells/ml with adipogenic differentiation was induced using MesenCult™ Adipogenic differentiation medium (Stem cell technologies). Following manufacturers protocol the stem cells were incubated with 2ml adipogenic differentiation medium and incubated at 37°C for 3 days or till lipid vacuoles observed under low magnification. Differentiated cells were washed with PBS, fixed with 10% formaldehyde for 15 minutes, dehydrated with 60% propanol, rinsed with water, air-dried, and stained with Oil Red O (Thermoscientific) dye diluted in 60% propanol for 10 minutes. Stained cells were imaged using a phase-contrast microscope (Olympus-BX53) with a 40X objective after mounting with DPX mounting media. Captured images were processed using cellSens software.

### 2.7 Chondrogenic Differentiation

MSCs were seeded on 22 mm coverslips at a density of 2 × 10□ cells/ml with 2 ml of MesenCult™-ACF Chondrogenic Differentiation Medium (Stem cell technologies) and incubated at 37°C and 5% CO2 for 3 days. On Day 6, and every 3 to 4 days thereafter, carefully removed the medium without disrupting the cells and replace it with 2mL of complete MesenCult™-ACF Chondrogenic Differentiation Medium till 21 days. Differentiated cells were washed with PBS, fixed with 4% formaldehyde for 15 minutes, washed with water, and incubated in 3% acetic acid for 5 minutes. Cells were then stained with Alcian Blue and Nuclear Fast Red solution diluted in 3% acetic acid for 60 minutes and washed with 1 M HCl for 5 minutes. The stained cells were imaged using a phase-contrast microscope (Olympus-BX53) with a 40X objective after mounting with DPX media. Captured images were processed using cellSens software.

### 2.8 Optimization of Secretome Collection

MSCs (1 × 10□ cells/ml) were cultured in a T25 flask until 70% confluence. The cells were washed twice with PBS, and DMEM without FBS but with antibiotics was added. The cells were incubated in a CO□ incubator for 24 and 48 hours. The media was collected and centrifuged twice at 2000 rpm for 10 minutes to remove cell debris. The supernatant was filtered through a 0.45 μm syringe filter, aliquoted, and stored at −80 °C. Similarly, HEK293 cells (1 × 10□ cells/ml) were cultured under the same conditions to collect normal cell-conditioned media as a control. The protein concentration was measured using Lowry method (14).

### 2.9 Effect of Secretome and Chemotherapeutic Drugs on Cancer Cell Lines (MTT Assay)

The cytotoxicity of chemotherapeutic drugs and stem cell secretome on cancer cell line MDAMB-231 was assessed using the MTT assay. Cancer cells (1 × 10□ cells/ml) were seeded in a 96-well culture plate and grown overnight in a CO□ incubator. Media were removed and replaced with PBS before incubation with serially diluted chemotherapeutic drugs (Cisplatin: 20 µg to 0.31 µg/ml; Paclitaxel: 40 µg to 0.625 µg/ml; Doxorubicin: 100 µg to 1.56 µg/ml; Vincristine: 200 µg to 3.12 µg/ml. Next, the secretome collected at 24 and 48 hours from ADSC, UCSC, and HEK293 were serially diluted (1000 µg/ml to 7 µg/ml) and incubated for 48 hours. After incubation, PBS washes were followed by the addition of MTT solution (yellow tetrazolium salt), which was reduced to purple formazan crystals. Propanol (100%) was added to dissolve the crystals, and absorbance was measured at 405 nm using a spectrophotometer. Cytotoxicity (%) was calculated after normalization with control samples.

#### 2.9.1 Mitochondrial Potential (TMRM)

HEK293, PC3, HepG2, and MDA-MB 231 cells were grown overnight and subsequently treated with stem cell secretome derived from adipose tissues and umbilical cords and HEK293 cells secretome as control. After 72 hours, cytotoxic effects were visualized under a fluorescent microscope by observing the mitochondrial membrane potential using TMRM dye. Treated cells were washed three times with 1x PBS, followed by the addition of 20 nM TMRM. The cells were incubated with TMRM for 45 minutes in the dark at room temperature. After incubation, the stained cells were washed three times with PBS and mounted with DPX for fluorescence microscopy. Images were captured at 573 nm using an Olympus (BX53) fluorescent microscope at 40x magnification. Captured images were processed using cellSens software.

#### 2.9.2 Effect of Secretome on Cell Migration (Scratch Assay)

MDA-MB 231 cells were grown overnight, and once they reached 70-80% confluency, a scratch was made using a 1 mm pipette tip in a straight line. The tip was held perpendicular to the bottom of the well, and light pressure was applied to remove the cell layer without damaging the plate. After scratching, the cell monolayer was gently washed to remove detached cells, and the wells were replenished with secretomes collected from stem cells and HEK293 cells (used as the control secretome). Scratched wells were imaged at 24 and 48 hours using a phase-contrast microscope (Olympus CKX41) at 4x and 10x magnification. Specific locations of the images were documented to ensure consistent imaging of the same spots over time. Subsequently the wound area created by the Scarth manually by tracing the cell free area within the captured image using Image J software.

#### 2.9.3 Profiling of Secretome

The stem cell secretome was precipitated, and the peptides were resuspended in a resuspension buffer (2% acetonitrile, 0.1% trifluoroacetic acid, and 0.1% formic acid in Milli-Q water). A volume corresponding to ∼1 µg of peptide was analyzed by nanoLC-MS/MS (Thermo Scientific Dionex Ultimate 3000 RSLC) coupled in-line with a HF mass spectrometer (Thermo Scientific Q Exactive) located at Sandor protieomics, Hyderabad, following their standard protocol. Raw data were searched against *Homo sapiens* using MaxQuant search engines and other protein databases. Only proteins with at least two unique peptide sequences, FDR <1%, and identified in all technical replicates (including the control, secretome from HEK293) were included. A difference in intensity of the same protein under different conditions of >3× was considered significant. The R statistical software version 3.3.2 (https://www.r-project.org) was used for further data analysis.

#### 2.9.4. Protein protein interaction and Mechanism of Action

The identified proteins were analyzed for protein-protein interactions and Gene Ontology (GO) biological processes. Using the online database STRING-DB (version 10.5), these proteins were categorized into functional groups, including tumor suppressors, apoptosis inducers, and DNA damage regulators.

##### Statistical analysis

The results are displayed as percentage of mean ± standard deviation (SD), as specified in the figure legends. Data analysis involved the use of appropriate statistical methods, to calculate the significance level and p value. All statistical analyses were with a significance level set at p<0.05, indicating statistical significance.

## 3. Results

### 3.1 Stem cell culture, lineage differentiation and characterization

Mesenchymal stem cells (MSCs) isolated from adipose tissue and umbilical cord tissue were cultured, and their morphological evaluation was performed using Giemsa staining. The spindle-shaped morphology confirms the fibroblastic nature of the stem cells (Suppl. Fig. 1A). The doubling times were determined as 75.71 hours for ADSCs and 50.03 hours for UCSCs (Suppl. Fig. 1B) suggests stable and robust proliferative potential ideal conditions for maintaining and expanding stem cells efficiently. Characterization of stem cells was performed using flow cytometry, with positive markers CD90, CD73, and CD105, and negative markers CD34, CD45, and HLA-DR. Both ADSCs and UCSCs showed strong positivity for CD105 (74–78%), CD73 (83–88%), and CD90 (94–96%) compared to negative markers (3.7–4.2%). Similar results were obtained through immunofluorescence, which are presented in the boxed sections of the flow cytometry diagrams for both ADSC and UCSC panels (Suppl. Fig. 1C). Stem cells were subjected to chondrogenic and adipogenic lineage differentiation using respective differentiation media. Alcian blue staining confirmed aggrecan expression in chondrocytes, while Oil Red O staining indicated lipid accumulation in adipocytes (Suppl. Fig. 1D).

### 3.1: Optimization of secretome collection and protein profiling

MSCs derived from adipose tissue and umbilical cord were cultured up to passage four (P4) and used for secretome collection at 24 and 48 hours. Passage 2 yielded a higher concentration of proteins compared to passages 3 and 4 (Suppl. Fig. 2A). Protein profiling revealed that the majority of proteins had a molecular weight of 50–150 kDa, followed by 150–250 kDa and 10–50 kDa (Suppl. Fig. 2B).

### 3.2: Optimization of cytotoxic effect of secretome at different passage with different time points

Secretomes collected at different passages were tested for their cytotoxic effects on MDA-MB-231 cells grown overnight and incubated for 24 and 48 hours. The UCSC-derived secretome from passage 2 (P2) exhibited a significant impact on cell viability compared to other passages (P3 and P4), as assessed by the MTT assay. In contrast, the ADSC-derived secretome followed a similar pattern to the UCSC-derived secretome but was less effective. Moreover, cell viability was more significantly reduced when MDA-MB-231 cells were exposed to secretomes derived from both ADSC and UCSC for 48 hours (Fig. 1A). To further validate cytotoxicity, mitochondrial membrane potential was assessed using TMRM (Tetramethylrhodamine) staining in cells treated with ADSC- and UCSC-derived secretomes for 48 hours. A decreased number of live mitochondria were observed in MDA-MB-231 cells treated with secretomes compared to control HEK293 cells (Fig. 1B).

**Figure 1:**
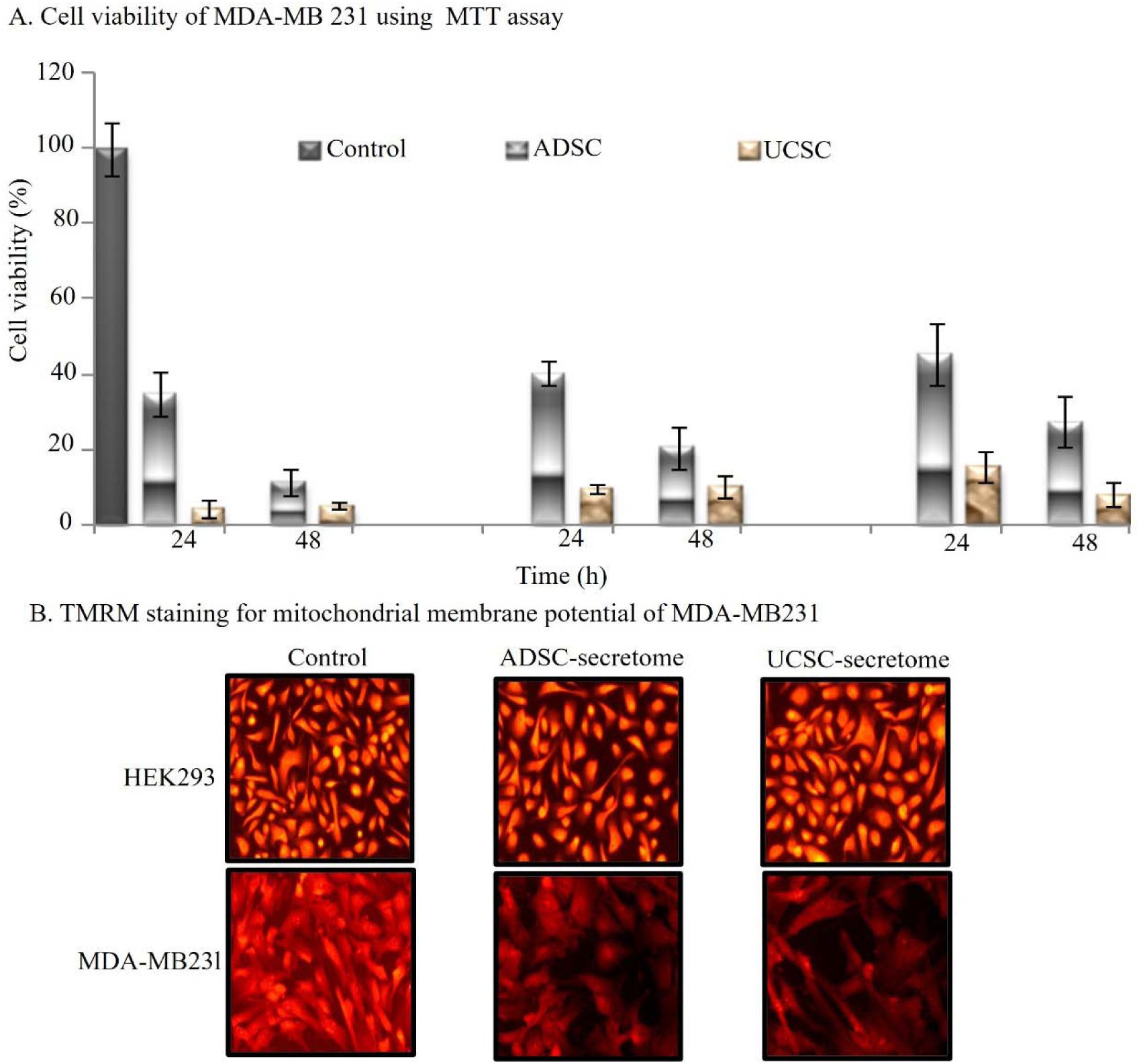
Stem cell secretome and optimization of cell cytotoxicity. A. MTT assay to check viability of MDA-MB 231 when incubated with stem cell secretomes collected at 24h and 48h of culture duration. HEK 293 cells secretome were used as a control. All the experiments were performed in triplicate for significant p-value (>0.5). B. Tetramethylrhodamine (TMRM) staining for mitochondrial membrane potential to the stem cell secretome treated MDA-mb 231 cells. Brighter red indicates healthy cells whereas dull red color with less cells indicates disrupted mitochondrial membrane. The image was taken fluorescent microscope with 20x objective after fixing with formaldehyde.

### 3.3: Stem cell secretome and cell cytotoxicity of MDA-MB231 cells

A cell scratch assay was performed to evaluate the effect of the stem cell secretome collected at passage 2 on MDA-MB-231 cell migration. It was observed that cells treated with the stem cell secretome exhibited a significant reduction in migration as early as 24 hours, with almost complete inhibition by 48 hours, compared to untreated cells (Fig. 2A). The time-dependent inhibition of cell migration is illustrated in the line diagram (Fig. 2B).

**Figure 2:**
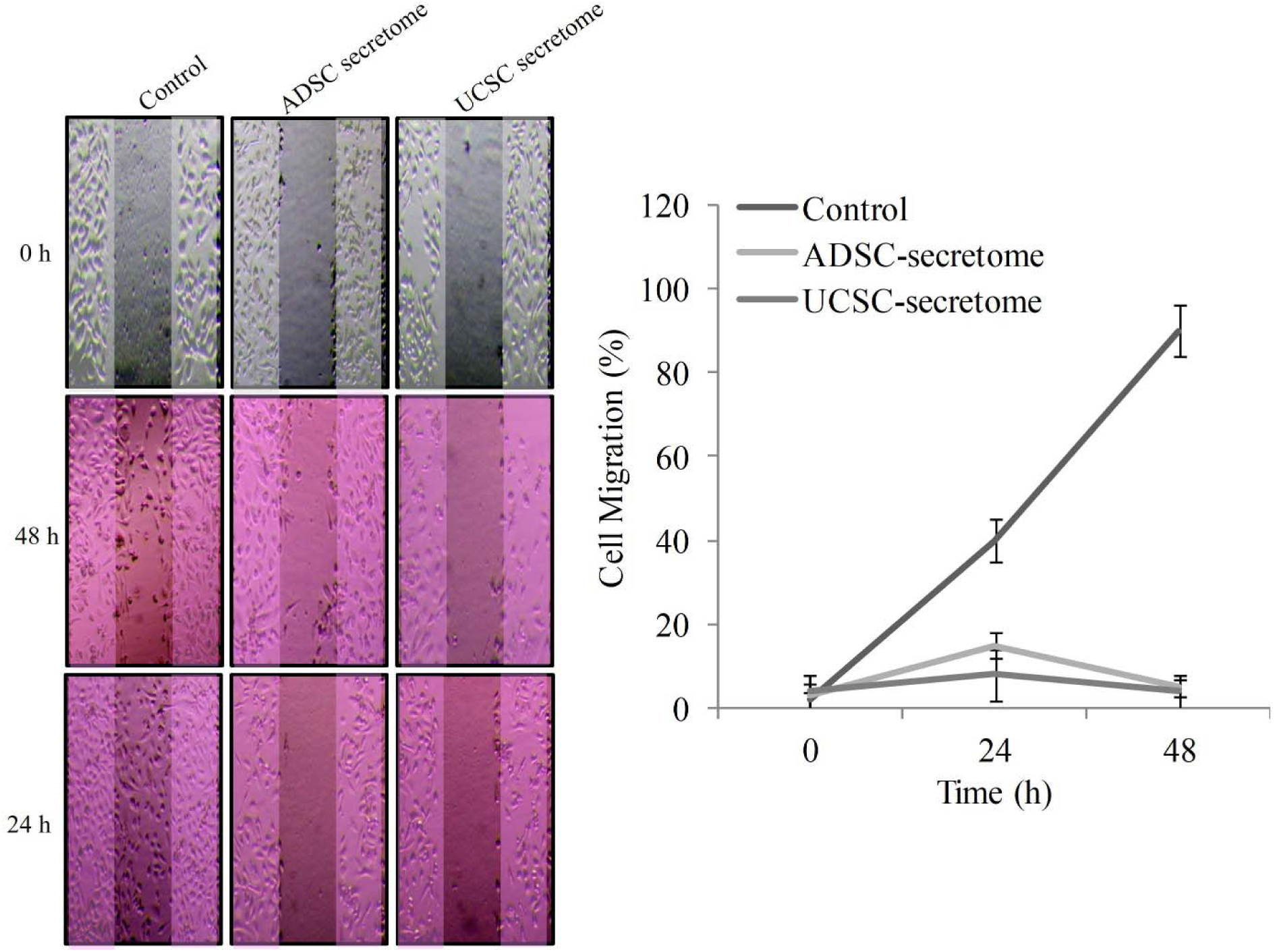
Effect of stem cell secretome on cell migration. Cell scratch on overnight grown MDA-MB 231 cell. Monitored cell migration for 24 h and 48h after the exposure of ADSC and UCSC stem cell secretome. Bright field microscopy at 4x objective was done to capture the cell movement at two different time points. Untreated cells were considered as control. Cell migration (%) was calculated using initial cell number and cell number after the secretome exposure (ADSC& UCSC) at two time points shown in line diagram.

### 3.4: Dose dependent anti-tumor effect of stem cell secretome on MDA-MB 231 cells

To evaluate the collateral damage of chemotherapeutic drugs (Doxorubicin, Cisplatin, Paclitaxel, and Vincristine) on normal cells, their cytotoxic effects were tested on MDA-MB-231 breast cancer cell line and HEK293 cells in a dose-dependent manner. It was observed that all the drugs exhibited cytotoxic effects on HEK293 cells (20%–40%), while the maximum tumor cell death achieved was only 50%–80%, even at the highest drug concentrations (Suppl. Fig. 3). To optimize the cytotoxic effect of stem cell secretome on MDA-MB-231 cells, secretome derived from P2-passaged ADSC and UCSC secretomes were tested at concentrations ranging from 7 ug/ml to 1000 ug/ml. MDA-MB-231 cells were incubated with these secretomes for 48 hours following overnight seeding. The MTT assay revealed a significant increase in cell death with increasing secretome concentration, ranging from 50% to 95% for UCSC secretome and 20% to 80% for ADSC secretome. In contrast, the secretome collected from HEK293 cells (control) exhibited minimal cytotoxic effects (<5% cell death) (Fig. 3).

**Figure 3:**
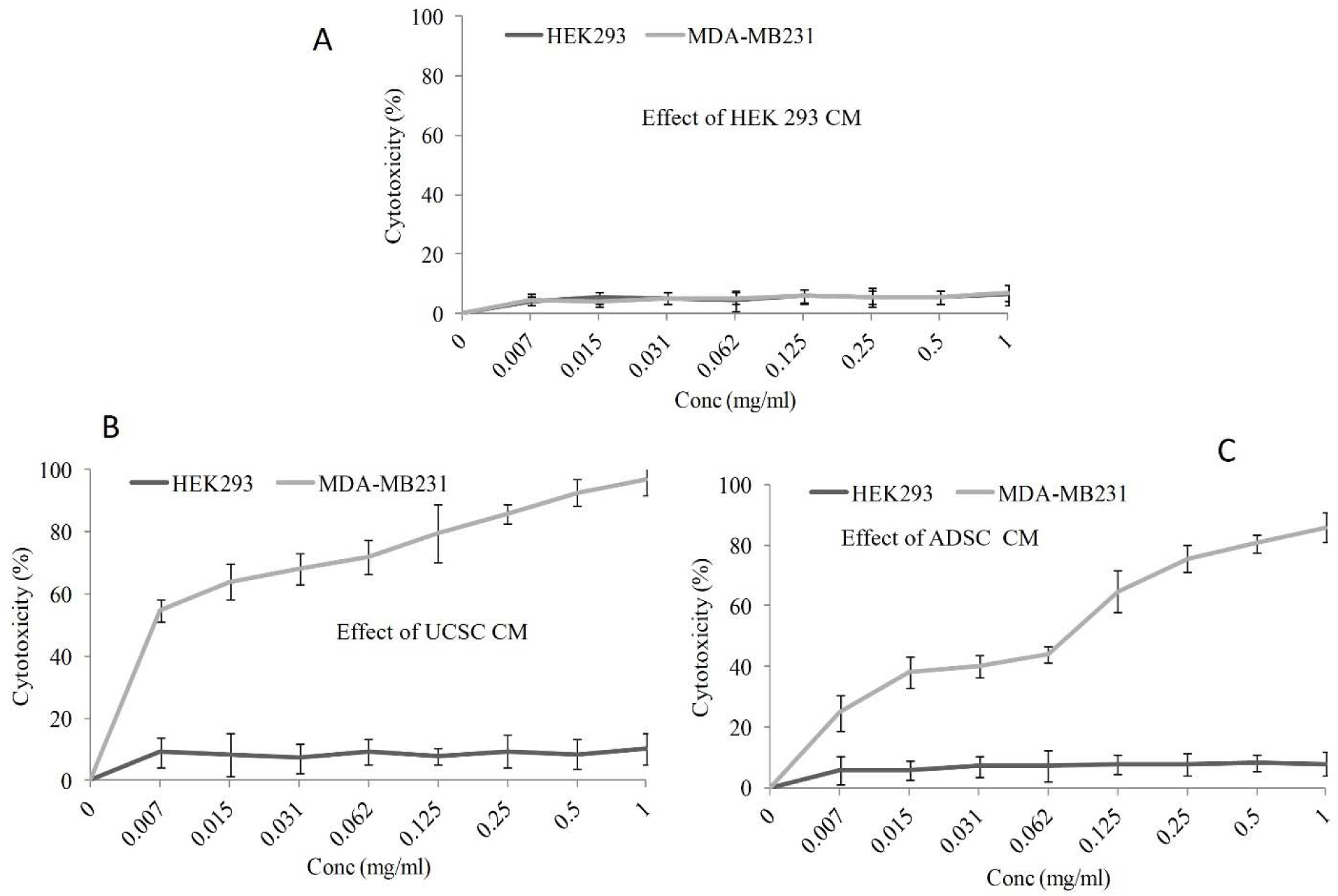
Anti-tumor effect of stem cell secretome on MDA-MB 231 cells. Serially diluted stem cell and HEK 293 cells (control) secretomes (1mg/ml to 0.007mg/l) were exposed to MDA-MB 231 cells for 48h. MTT assay exhibits the cell cytotocity (%) of MDA-MB 231 in line diagram. All the experiments were performed in triplicate to achieve highly significant data.

### 3.5: Protein interaction and networking of secretome proteins

LC-MS protein profiling data of ADSC and UCSC secretomes revealed the presence of unique secretory proteins in both types of secretomes. The UCSC secretome exhibited sharper peaks between 37 and 44 minutes, whereas the ADSC secretome showed broader peaks appearing slightly later, between 39 and 45 minutes. Additionally, an extra peak in the ADSC secretome was observed between 20 and 25 minutes, which was absent in the UCSC secretome. The control secretome (HEK293 cells) displayed peaks only between 1 and 9 minutes. The ADSC secretome exhibited unique peaks with lower intensity, which were absent in both the UCSC secretome and the control (Suppl. Fig. 4). Profiling of stem cell secretomes derived from ADSC and UCSC identified various proteins with tumor-suppressing properties, apoptosis-inducing functions, DNA damage response mechanisms, and other associated functions that contribute to tumor growth inhibition. Using STRING (a functional protein association network), the interactions among these proteins were mapped to demonstrate their protein-protein interactions and potential mechanisms of action in suppressing tumor progression (Fig. 4).

**Figure 4:**
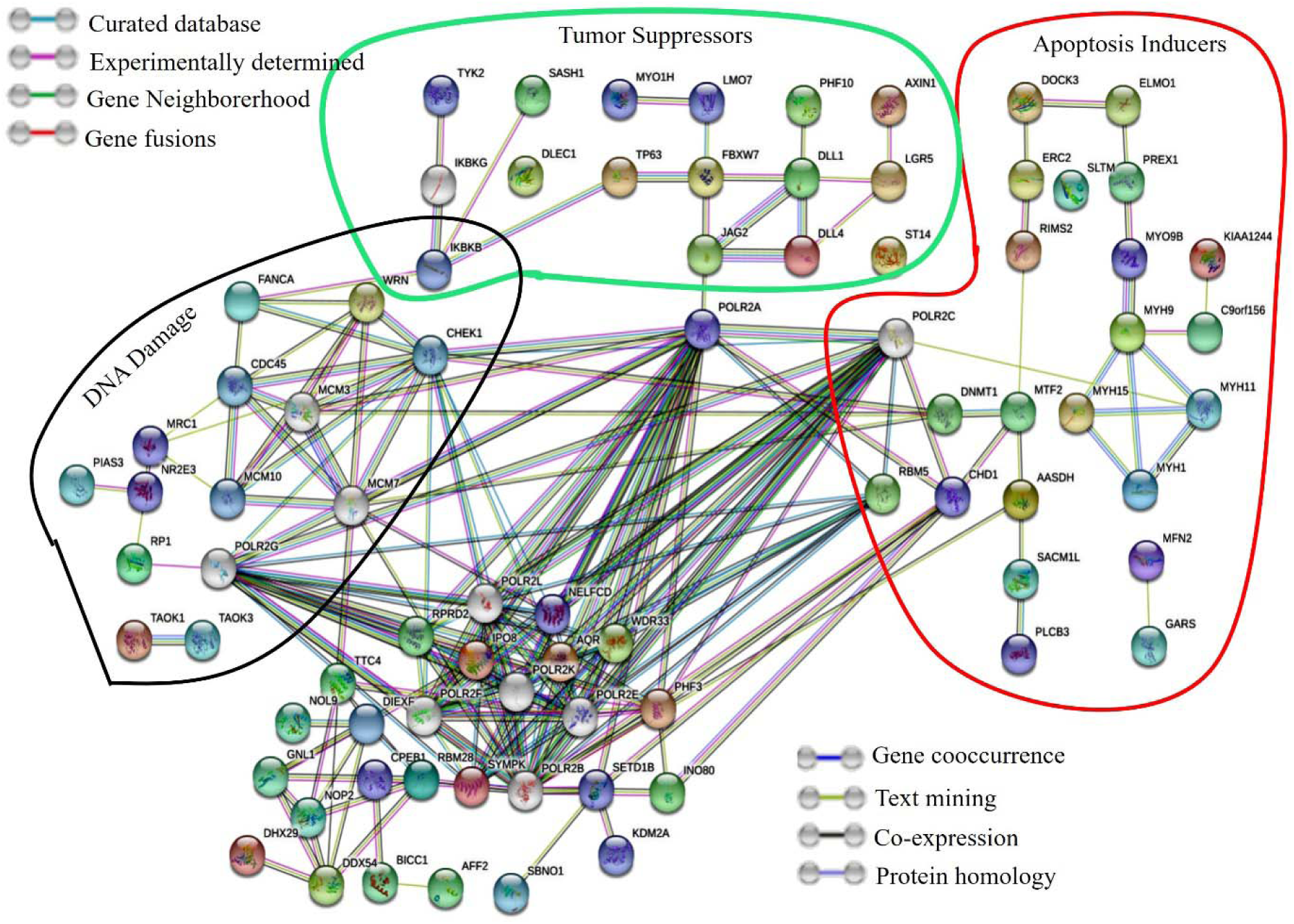
Protein-Protein interactions and networking of profiled proteins using STIRNG. Significant proteins obtained from the LCMS profiling of secretomes were submitted to STRING (https://string-db.org/) for possible interactions and enrichment analysis. Red circles indicate apoptotic proteins, green circles indicate tumour suppressors and black circles indicates the proteins which are involved in DNA damage. These proteins with blue lines were from curated database, pink lines were experimentally verified by others, green lines indicate gene neighbourhood, red line for gene fusions, blue line indicates gene co-occurrence, light green lines mean text mined genes etc.

## Discussion

With decades of research, it has become evident that chemotherapeutic drugs induce long-term side effects, which pose inevitable challenges for both clinicians and patients. Despite several advancements in therapeutic interventions, the efficacy and side effects of chemotherapy have not significantly improved, remaining a major concern for enhancing cancer patient survival and reducing financial burdens. Therefore, developing new approaches for cancer management is imperative to improve therapeutic efficacy while minimizing the side effects of chemotherapy. A promising strategy involves treating cancer patients by promoting repair mechanisms, restoring diseased tissues, and rejuvenating quality of life through the use of stem cells or their derivatives (15). The mesenchymal stem cell (MSC) secretome plays a crucial role in modulating the tumor microenvironment, influencing cancer growth dynamics, and harnessing rejuvenation potential for promising therapeutic outcomes. Research findings collectively indicate that stem cells and their derivatives hold significant potential for cancer therapeutics and regeneration (16). In this study, we demonstrate how leftover tissues, such as adipose tissue and umbilical cord, which are typically discarded after surgery, can serve as a valuable source of mesenchymal stem cells (MSCs) for cancer therapeutic interventions. Stem cells possess unique properties, including immune modulation, selective migration to inflammatory sites, and secretion of growth factors, with therapeutic potential demonstrated in prostate cancer (17).

A well-established and characterized stem cell population (ADSC & UCSC) was optimized, and the secretome was collected for evaluation of its anti-cancer properties on breast cancer cell lines (MDA-MB231) in vitro. The umbilical cord, which is typically discarded after childbirth, has recently emerged as an excellent source of mesenchymal stem cells. Cytotoxicity assays reveal that breast cancer cell migration and survival are significantly impaired when incubated with stem cell secretomes, indicating that key secretome components block cancer progression and initiate cell death by expressing apoptotic inducers, tumor suppressors, and DNA damage regulators, as observed in the protein-protein interaction network (18). The anti-tumor effects of stem cell secretomes include blocking signaling pathways involved in proliferation and cell cycle regulation, ultimately reducing tumor growth (19). Notably, umbilical cord-derived stem cell secretomes significantly decrease cell viability and proliferation by inhibiting PI3K/AKT activation (20).

A dose-response cytotoxicity assay was performed to evaluate the impact of secretomes on cancerous (MDA-MB231) and normal (HEK293) cells. The results demonstrate a selective cytotoxic effect on cancer cells while sparing normal cells, suggesting that stem cell-derived secretomes can act as tumor suppressors without causing collateral damage to healthy tissues. This finding is particularly relevant for personalized medicine and targeted cancer therapy, where minimizing side effects is a top priority (1).

To further elucidate the molecular pathways influenced by stem cell secretomes, gene enrichment and network analysis were performed. The analysis categorizes genes into functionally significant groups, including tumour suppressors, apoptosis inducers, and DNA damage regulators. The protein interaction network highlights the presence of key regulatory proteins that inhibit tumour progression, including genes involved in cell cycle arrest, DNA repair, and immune modulation, which collectively contribute to cancer suppression (22). The apoptosis induction network (red region) indicates that secretomes contain molecular signatures that promote programmed cell death in cancer cells. The activation of apoptotic pathways is crucial for eliminating malignant cells while preventing metastasis.

Stem cell secretomes represents a novel and innovative approach in modern medicine. Unlike conventional cell-based therapies, which face challenges such as regulatory hurdles, ethical concerns, and risks like immune rejection, secretome-based therapy bypasses these limitations while retaining the therapeutic benefits of stem cells. The bioactive components of secretomes act as molecular mediators that influence cell fate, making them an ideal candidate for regenerative medicine and cancer treatment.

## Conclusion

Stem cell-derived secretomes demonstrate an innovative approach in cancer managemnet paving away for targeted, non-invasive, and highly potent alternative to conventional therapies. By harnessing the natural regenerative and immune-modulating properties of stem cells, secretomes show immense promise in treating aggressive cancers, such as breast cancer. This study underscores the potential of secretomes derived from stem cells from surgical waste as a novel and non-invasive cancer therapy. The findings suggest that these secretomes contain bioactive molecules capable of selectively targeting cancer cells while preserving normal cellular functions, making them a highly attractive candidate for future cancer treatments. As per vitro finding, further research is needed to explore their therapeutic potential in vivo models and, ultimately, in human clinical trials. Future studies should focus on optimizing secretome formulations, identifying key bioactive components, and assessing their clinical efficacy to advance this revolutionary therapy towards real-world applications.

## Conflict of Interest

There are no conflicts of interest.

## Acknowledgement

We would like to express our gratitude to Mr. Mohammed Abdul Razak for his invaluable assistance in gathering patient information samples and initial experiments. Additionally, Mr. Prasad provided vital support with flow cytometry instrumentation and data management.

## Funding

KIMS foundation and Research Center (KFRC)

## Approval

Ethics Committee, KIMS Hospitals (KIMS/EC/2020/66-0), Minister Road, Secunderabad, Hyderabad, Telangana, India.

## Authors Contribution

The conceptualization, experimental design and data analysis was conducted by VKV and SSB, who also authored the original draft. NKP and KS were responsible for conducting the experiments and compiling data. RC provided instrumentation support, facilitates the samples questions and corrections of final draft.

## Figure Legend

**Supplementary2 Figure 1:**
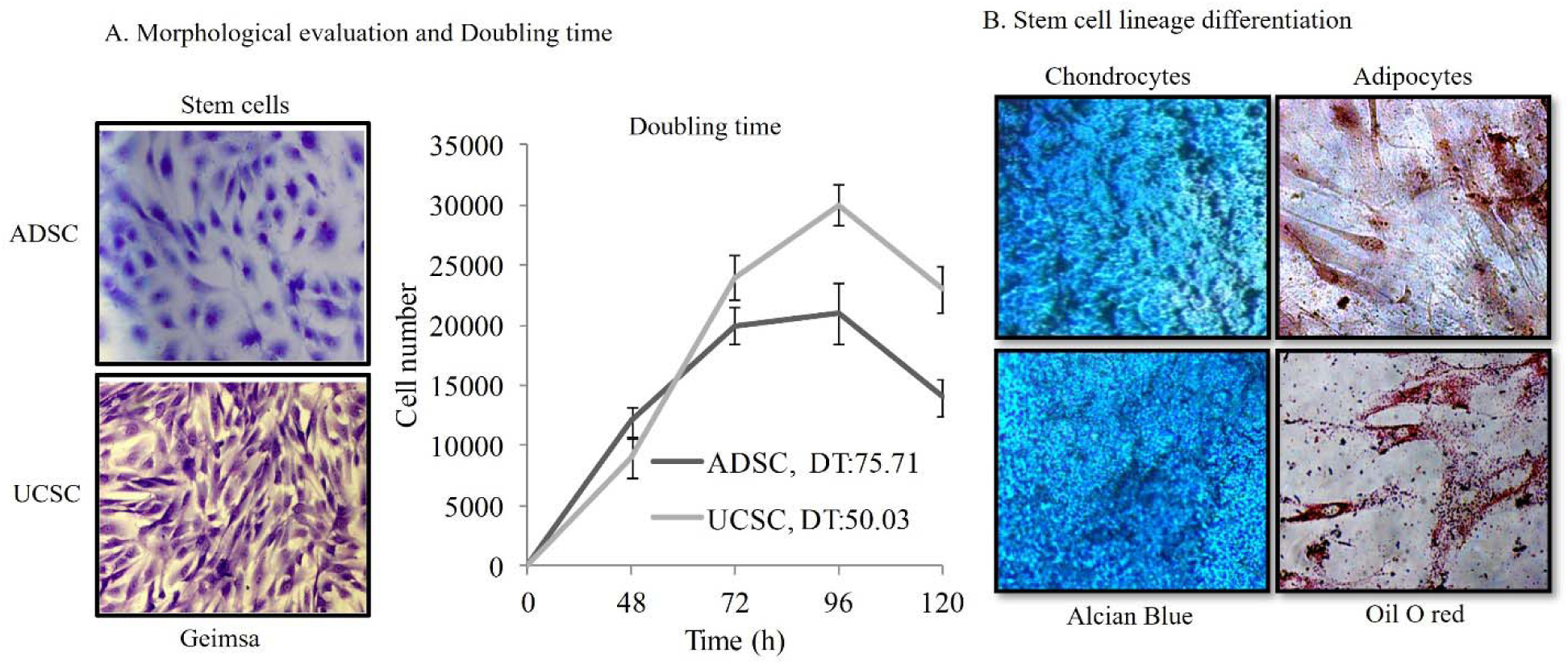
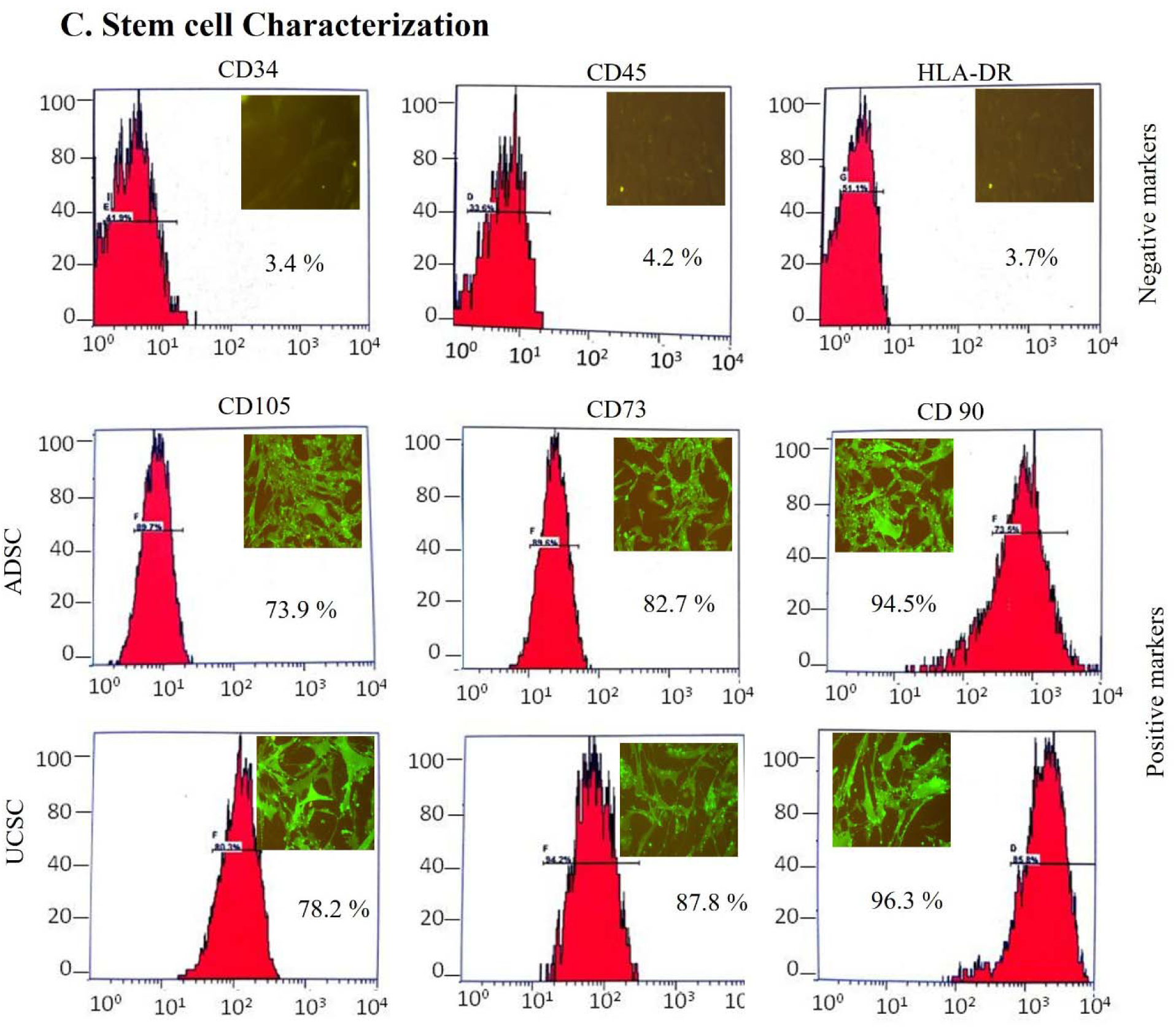
Stem cell culture, lineage differentiation and characterization. A. Giemsa staining (blue color) to show morphology of ADSC and USCS. B. Doubling time of ADSC and UCSC calculated using formula online with triplicate experiments for appropriate significance. C. Characterization of ADSC and UCSC using monoclonal antibody of positive markers (CD105, CD73, CD90) linked with Alexa-488 and negative markers (CD34, CD45, HLA-DR) linked with PE was done on flow cytometry. IgG isotype control was used to compensate and gating. Similar antibodies were used for immunofluorescence of positive and negative markers on the ADSC and UCSC grown on cover slips. 40X image of all marker’s expressions are shown in green color inside the flow cytometer histogram plot. D. Adipogenic and chondrogenic differentiation staining with Chondrocytes with alcian blue (blue color) and adipogenic with Oil-red O (red color).

**Supplementary Figure 2:**
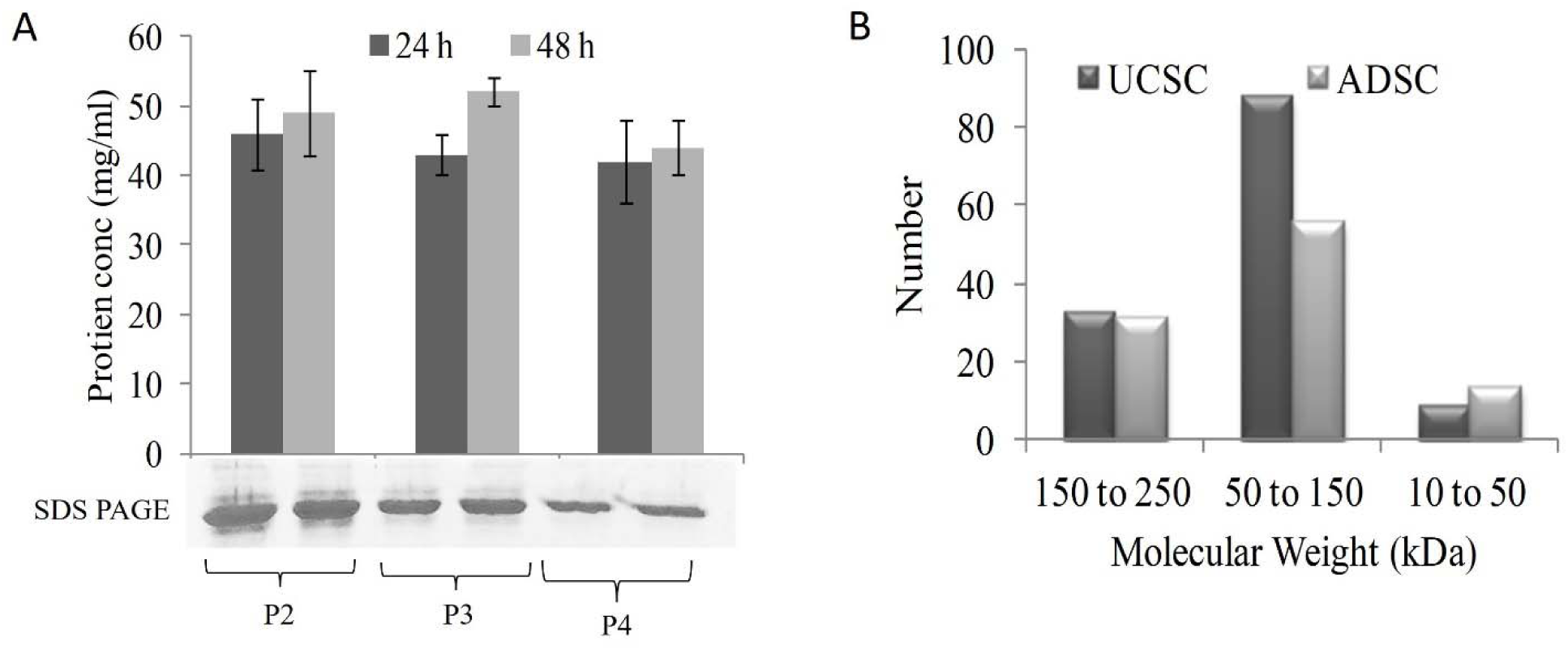
Collection of secretomes from stem cells. A. SDS-PAGE represents protein concentration (mg/ml) of stem cell secretomes from passage (P2, P3 & P4) at 24 h and 48h durations which are depicted as bar diagrams. B. Representation of number of proteins with respective molecular weight (KDa) obtained from protein profiling using LCMS form UCSC and ADSC stem cell secretomes. Experiments were performed in triplicates which are shown as error bar.

**Supplementary figure 3:**
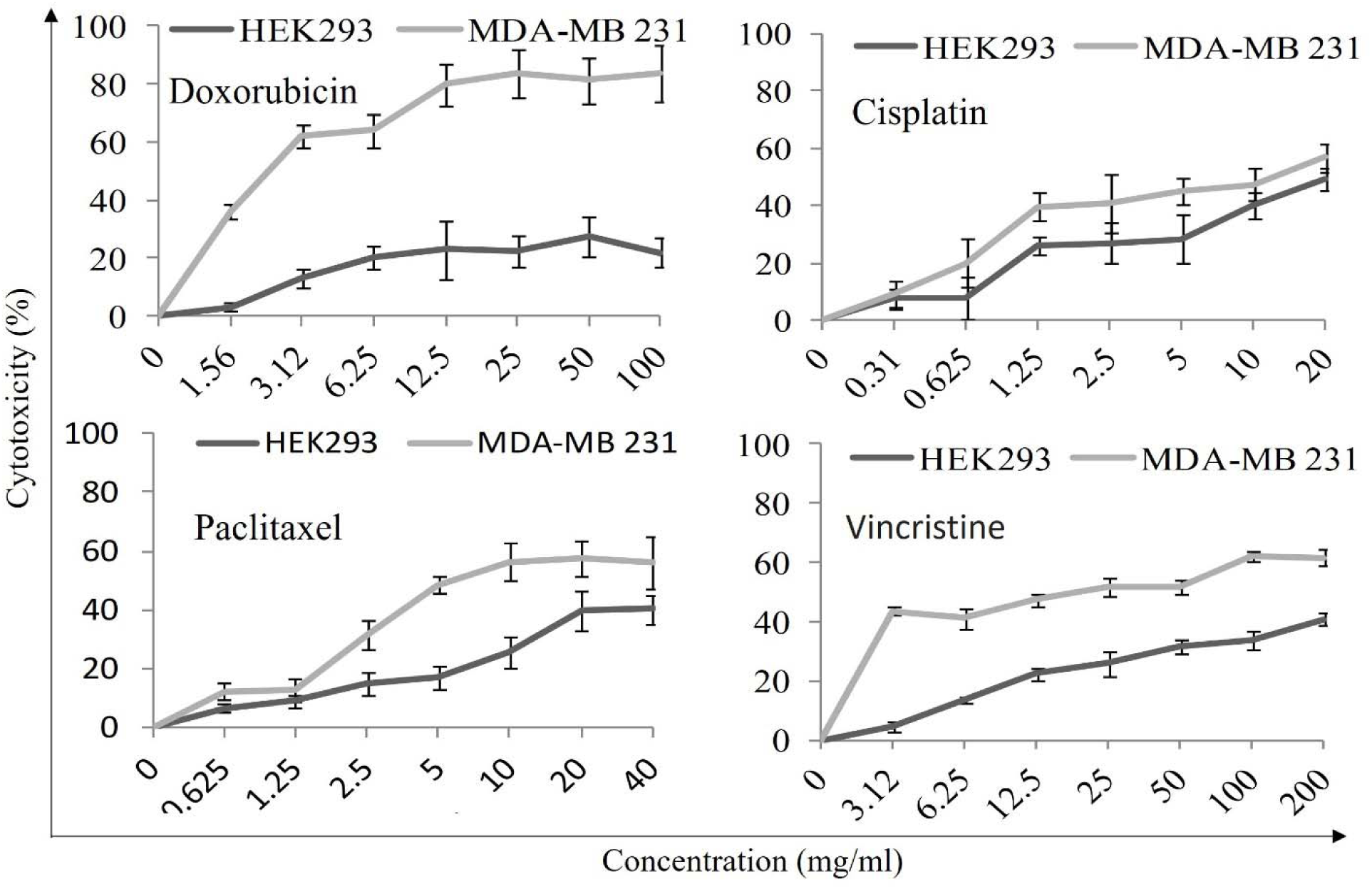
Effect of anticancer drugs (commercially available) on MDA-MB 231 cells. MTT assay represents the cytotoxic effect of Doxorubicin (1.56 to 100 mg/ml), Cisplatin (0.31 to 20 mg/ml), Paclitaxel (0.625 to 40 mg/ml) and Vincristine (3.12 to 200 mg/ml) concentrations serially diluted from highest to lowest concentration. Significant p value (> 0.05) obtained by repeating the experiments more than thrice.

**Supplementary figure 4:**
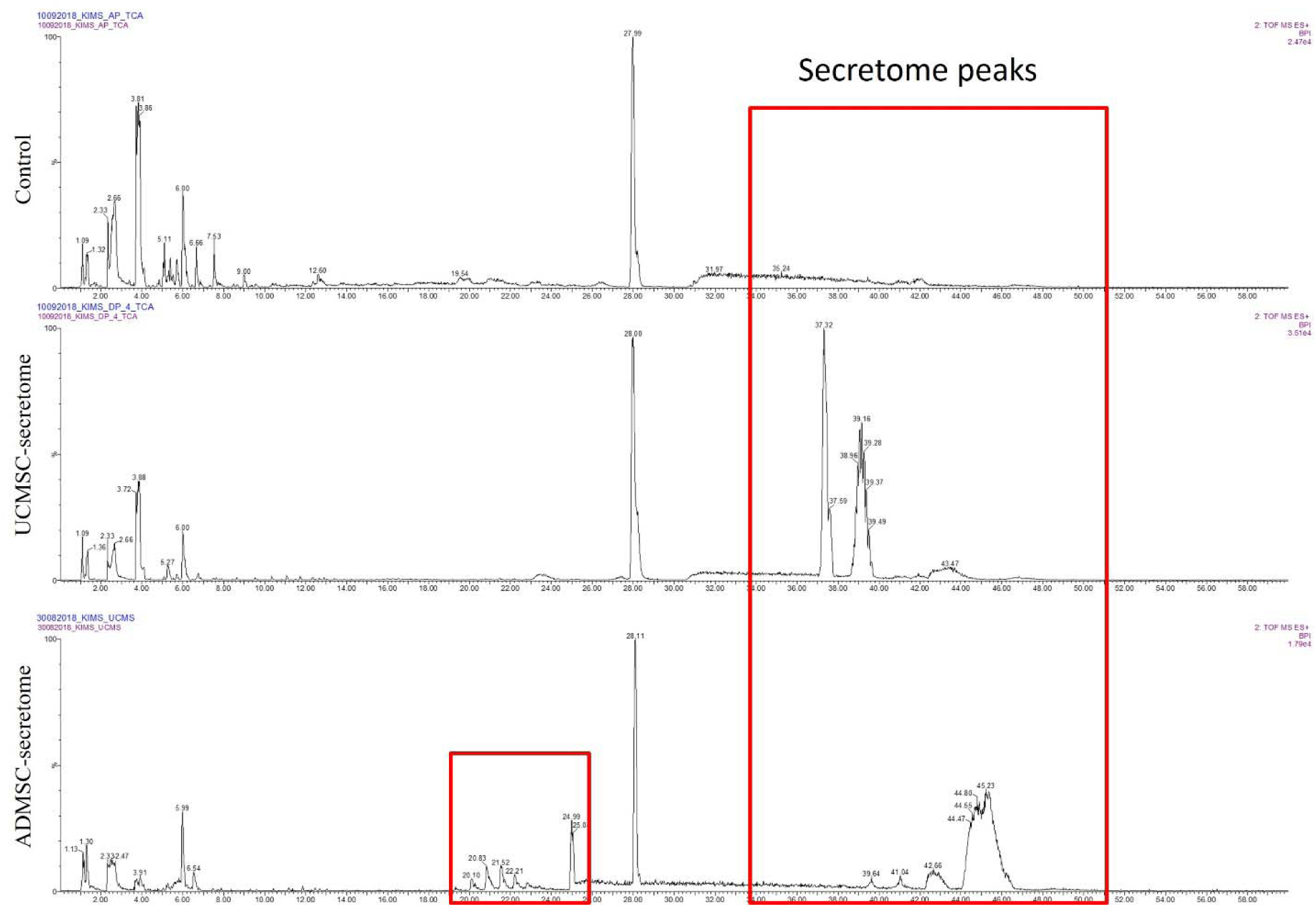
Profiling of stem secretomes using LCMS platform. LCMS protein profiling was outsourced from Sandor Proteomics, Hyderabad, whereby following the standard operating procedure of LCMS data was received as EI spectrum of peptides (%) at x-axis against retention time (0-58 Min) at y-axis. Experiment was done in triplicate to remove the false positive proteins and increase the significance level.

## Notes

### Competing Interest Statement

The authors have declared no competing interest.

