## Supplementary figures and images for "Stem Cell Secretomes from Surgical Waste: A Novel Approach to Cancer Therapy – An In Vitro Study"

### Suppl. Fig. 1A

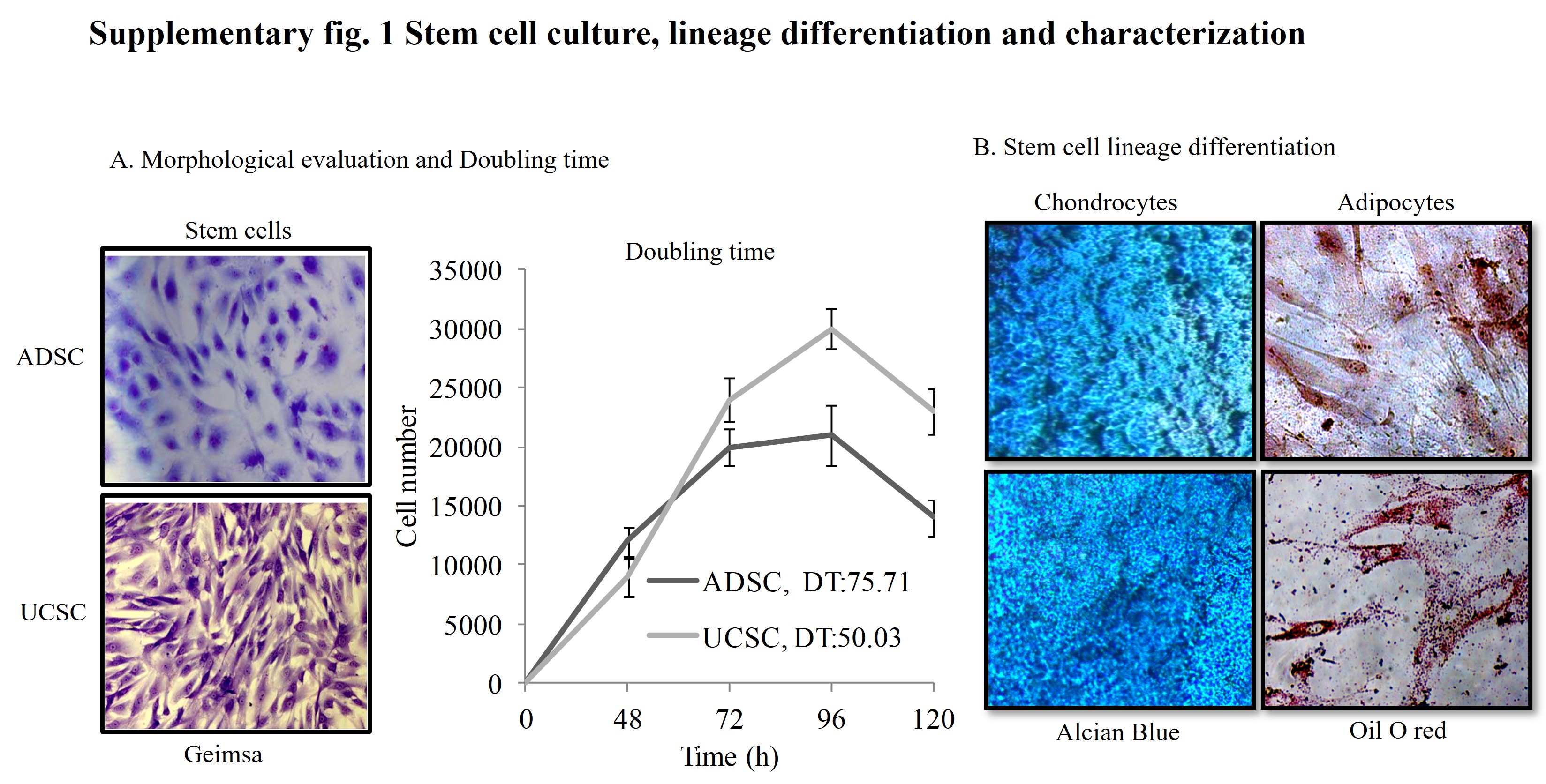

### Suppl. Fig. 1C

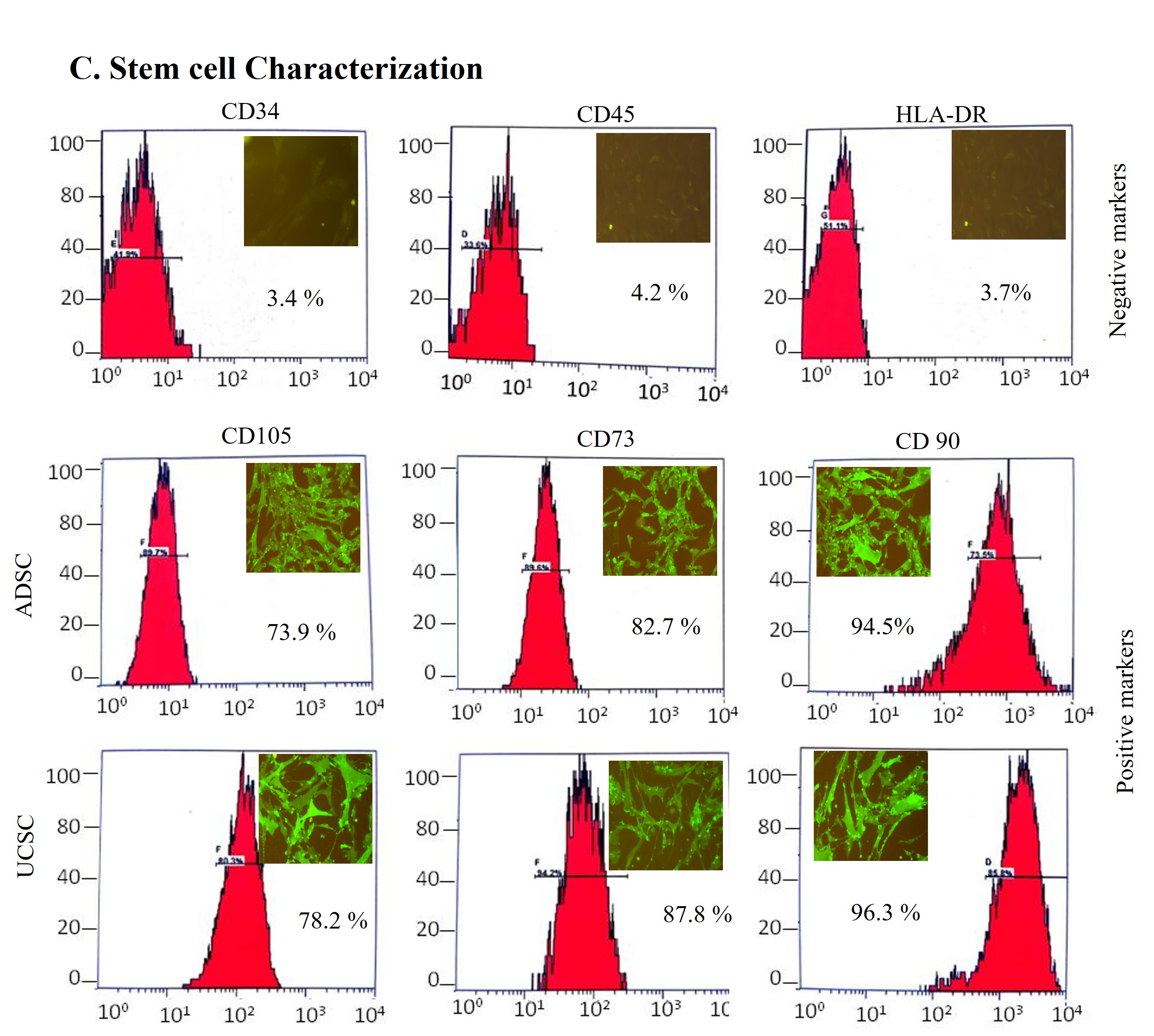

### Suppl. Fig. 2A

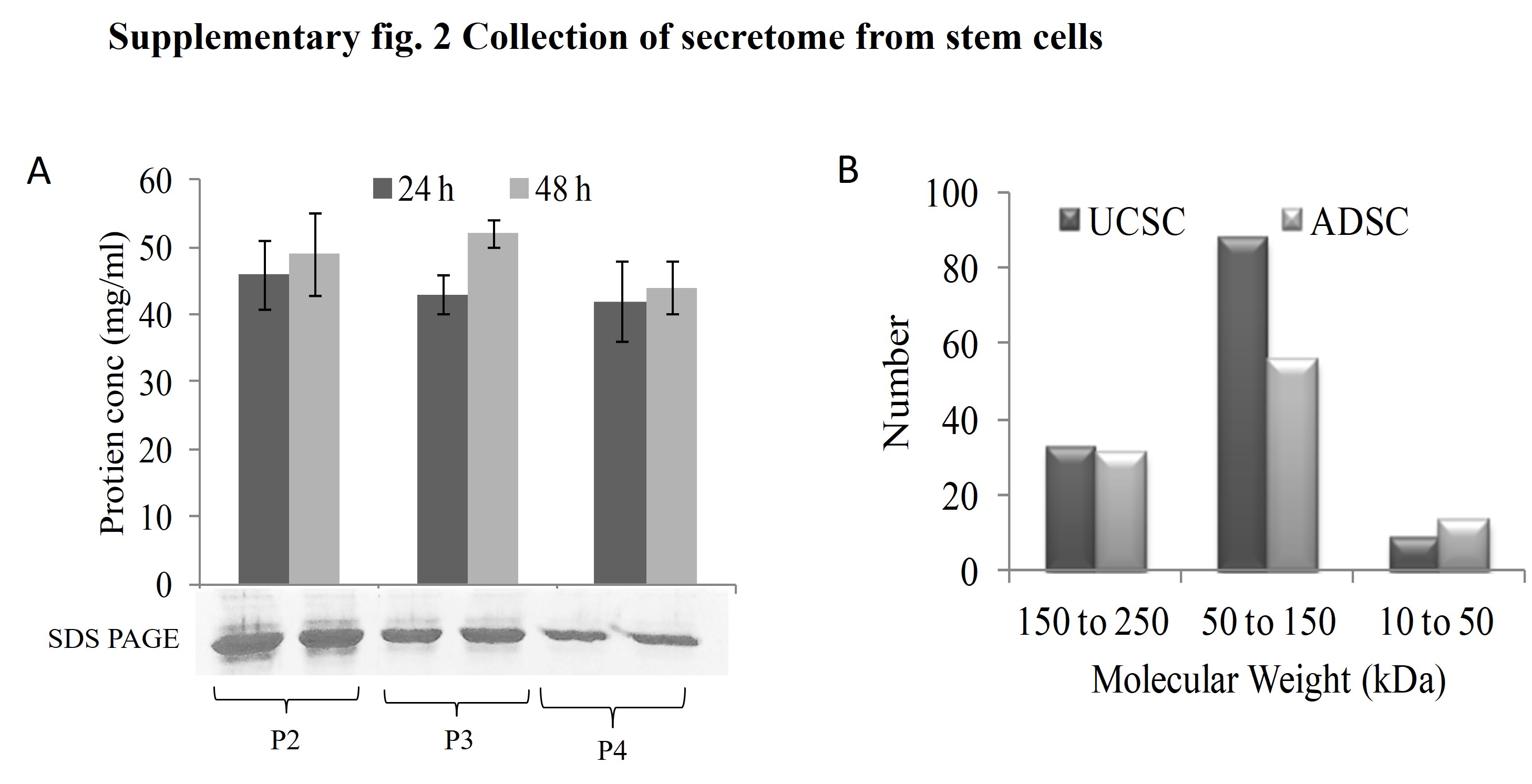

### Suppl. Fig. 3

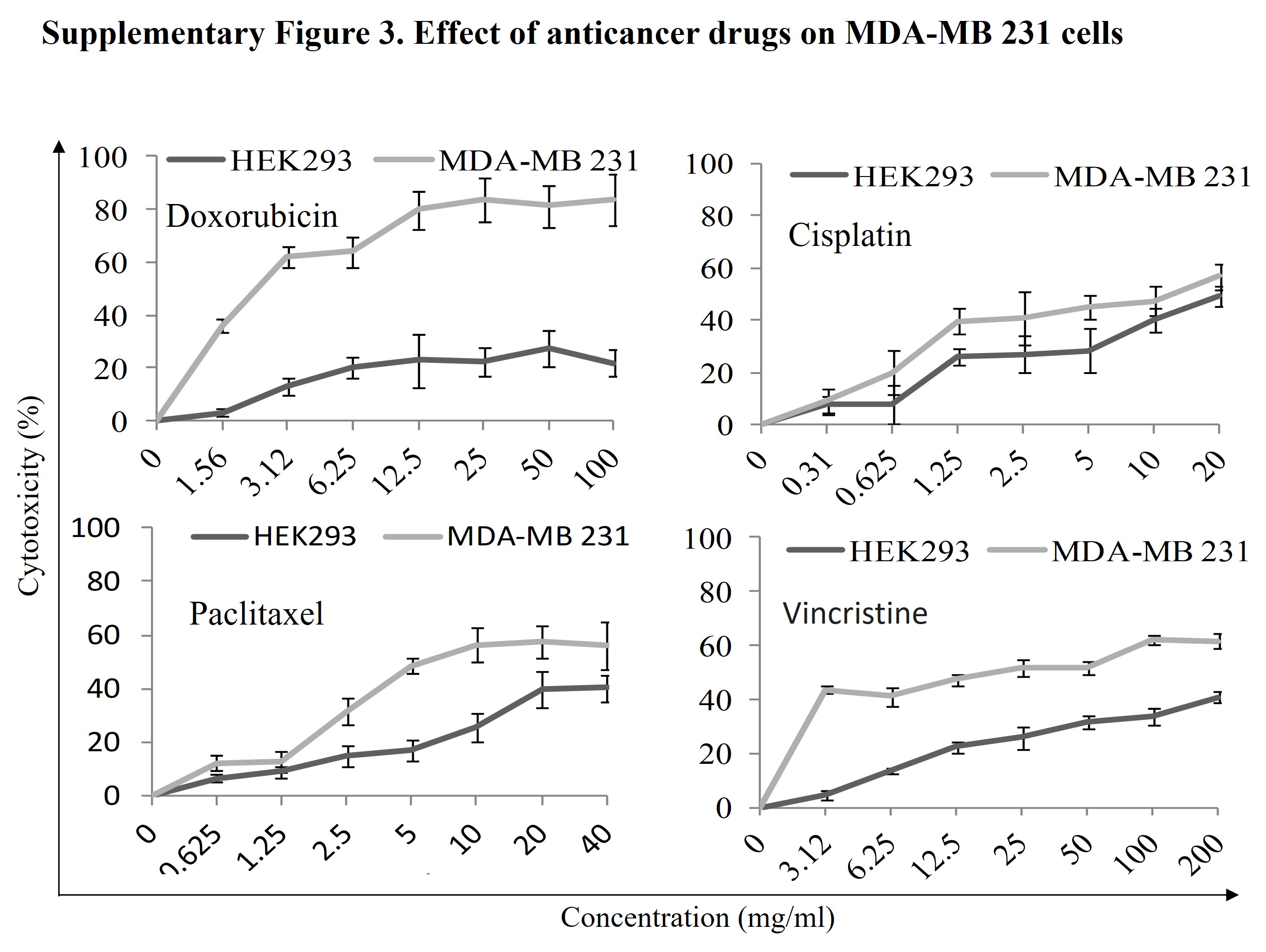

### Suppl. Fig. 4

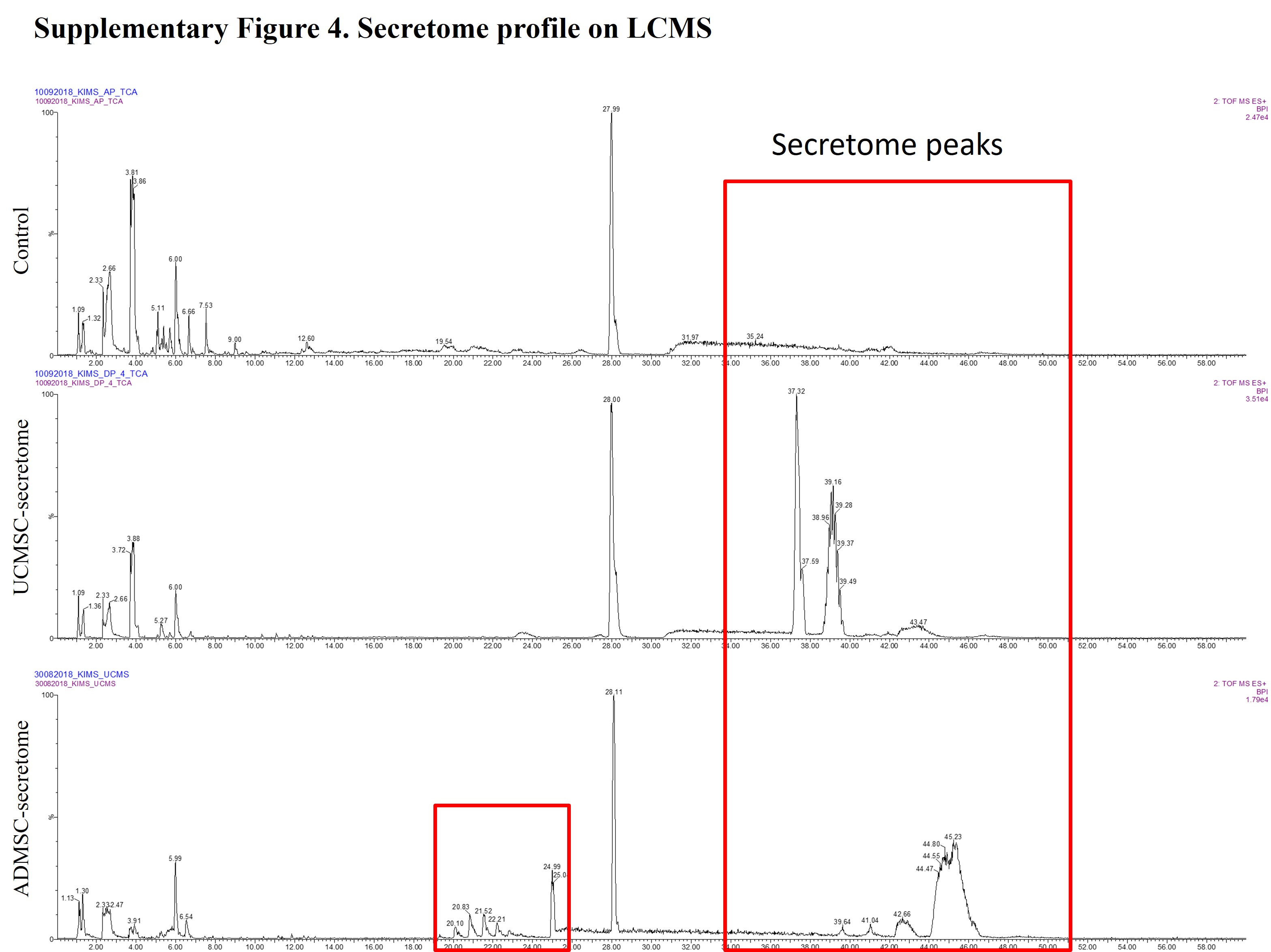
